# Targeting histone acetylation enables epigenetic modulation of inflammatory pathways – a novel therapeutic strategy for rheumatoid arthritis

**DOI:** 10.1101/2025.01.14.632975

**Authors:** Marie Brinkmann, Anela Tosevska, Bianca Luckerbauer, Moritz Madern, Lisa Göschl, Wilfried Ellmeier, Daniel Aletaha, Teresa Preglej, Michael Bonelli

## Abstract

Autoimmune diseases like rheumatoid arthritis (RA) are characterized by a systemic inflammation caused by autoreactive immune cells. Epigenomic modulation of these cells offers a strategy to reprogram pathogenic pathways without altering the genome, potentially restoring immune balance. Epigenetic inhibitors are already utilized in oncology but often exhibit adverse effects due to lack of selectivity and cytotoxic concentrations. Applying these drugs to treat autoimmune diseases necessitates more selective inhibitors and the use of tolerable concentrations. In this study, we screened a library of 25 compounds with varying degrees of target selectivity and different concentrations. Spectral cytometry enabled the analysis of cell-subset distribution and activation, followed by bulk RNA-sequencing for transcriptomic profiling.

We could demonstrate cell-subset specific and concentration-dependent immune modulation in PBMCs. Transcriptomic analysis showed that inhibitors of histone acetylation-modulating enzymes significantly altered gene expression, particularly in immune regulation pathways relevant to autoimmune diseases. Comparative analysis between *in-vitro* treated healthy controls and RA patients demonstrated both shared and selective drug effects, with some inhibitors like Ricolinostat overlapping with established RA drug pathways.

Our findings highlight the potential of epigenetic inhibitors, especially those targeting histone acetylation, to modulate immune responses in a target-selective manner.

**Graphical abstract:** 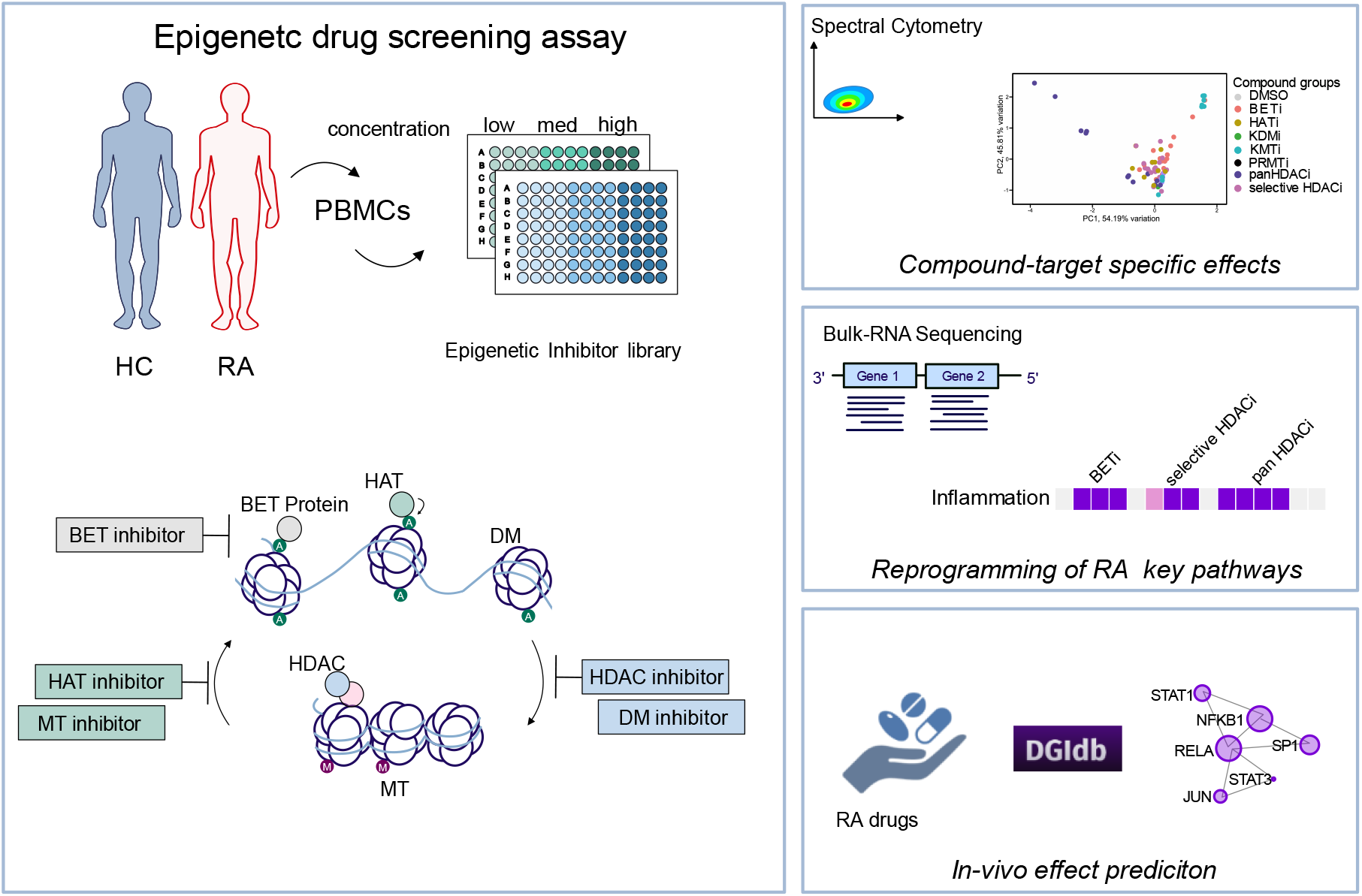

## Introduction

Rheumatoid arthritis (RA) is a chronic systemic autoimmune disease characterized by the presence of autoreactive immune cells and a loss of self-tolerance (1, 2). These pathogenic cells infiltrate the synovium, leading to painful joint swelling, bone resorption, and ultimately, disability and reduced quality of life (3).

Epigenetic regulators, including histone-modifying enzymes, have been implicated in multiple inflammatory conditions (4, 5). The interplay of these enzymes determines histone modifications, such as acetylation and methylation, which influence chromatin accessibility and gene expression. Histone acetylation is regulated by histone acetyltransferases (HATs) and histone deacetylases (HDACs), while bromodomain and extra-terminal (BET) proteins bind acetylated histone residues and act as further transcriptional regulators. Histone methylation is controlled by histone methyltransferases (MTs) and demethylases (DMs) (6). Beyond histones, these enzymes can also modify non-histone targets (7).

Histone-modifying enzymes play critical roles in inflammation and immune cell regulation. For instance, histone acetylation regulates the activity of transcription factors (TFs), including Nuclear Factor kappa B (NF-κB), a central driver of cytokine production, inflammatory signaling, and osteoclastogenesis in RA. (8-10). Acetylation modifying enzymes also affect cell fate and differentiation processes in immune cells central to RA pathogenesis, such as inflammatory T helper cells and monocytes/macrophages (9, 11, 12). Further, we have recently demonstrated that CD4^+^ T cell conditional knockout mice were protected from developing collagen-induced arthritis (CIA), underscoring the role of histone acetylation in inflammatory joint diseases (13). A dysregulation of histone methylation is known to promote autoimmune processes by enhancing pro-inflammatory gene expression and impairing regulatory mechanisms (14). Specifically, selective MTs such as Enhancer of zeste homolog 2 (EZH2) are found to be overactive in rheumatic diseases (15).

Targeting epigenetic modifiers has proven effective in cancer therapy. However, pan-HDAC inhibitors (pan-HDACi) have demonstrated considerable side effects in clinical trials, highlighting the need for more selective and less cytotoxic approaches in RA treatment (16). Preclinical studies have shown promising results for HDAC and BET inhibitors in reducing inflammation and joint swelling in murine RA models (17-19). Furthermore, the pan-HDACi Givinostat improved clinical outcomes in a phase II trial for juvenile arthritis (20). However, the precise mechanisms of these compounds on human immune cells, particularly in RA, remain poorly understood. Notably, HDAC inhibition is not directly translatable to knockout models due to the involvement of HDACs in multiprotein complexes, emphasizing the need for testing selective inhibitors.

In this study, we systematically screened an epigenetic inhibitor library targeting pan-HDACi, (class-) selective HDACi, HATi, BETi, lysine demethylases (KDMTs), lysine methyltransferases (KMTs), and arginine methyltransferases (PRMTs) across various concentrations. Our findings demonstrate compound– and target-selective effects, with shared modulation of pathways targeted by established RA therapies, underscoring the potential of epigenetic reprogramming as a novel therapeutic strategy.

## Methods

### Human subjects and ethical aspects

Blood samples from healthy individuals and RA patients were collected at the Division of Rheumatology at the Medical University of Vienna. This study was carried out in accordance with the recommendations of the “guidelines for storing and using patient samples” of the Ethical Committee of the Medical University of Vienna. All subjects gave written informed consent in accordance with the Declaration of Helsinki that their blood can be stored and used for scientific purposes. The protocol was approved by the Ethical Committee of the Medical University of Vienna (ethics vote number: 2071/2020, 1075/2021).

### Cohort information

Blood from healthy controls (n=3) was used to screen 25 compounds at 3 different concentrations via flow cytometry. For bulk-RNA sequencing, PBMCs from an additional 3 healthy controls were utilized. A group of RA patients (n=3), matched for sex and age, was included. These patients were drug-free at the time of sample collection and showed no laboratory-detectable indicators of autoimmune disease.

### Blood Collection and PBMC Isolation

Blood samples were collected in heparinized blood collection tubes. Peripheral blood mononuclear cells (PBMCs) were isolated using Pancoll density gradient centrifugation. Briefly, 10 mL of whole blood was diluted 1:1 with PBS, carefully layered over 15 mL of Pancoll (PAN-Biotech) solution, and centrifuged at 530 g for 22 minutes at room temperature (RT) without a brake. Following centrifugation, the distinct white layer containing PBMCs was transferred into a new 50 mL Falcon tube, washed with PBS, and centrifuged at 400 g for 8 minutes at 4°C. The resulting cell pellets were resuspended in 5 mL of PBS. PBMCs were counted within a size range of 7–15 μm using a Z2 Coulter Particle Count and Size Analyzer (Beckman Coulter). After another spin (400 g, 8 minutes, 4°C), the PBMCs were resuspended in freezing medium (RPMI-1640 with 20% FCS [Gibco] and 15% DMSO [Sigma]) at a concentration of 10–20 × 10^6^ cells/mL, aliquoted into cryovials. and placed in a CoolCell FTS30 freezing container for 24 hours before being transferred to liquid nitrogen for long-term storage.

### Thawing of PBMCs

Cryovials containing 5– 10 ×10^6^ PBMCs were thawed by incubating them at 37°C for 5 minutes. The contents were then transferred into a 50 mL Falcon tube. Cell culture medium (RPMI-1640 supplemented with 10% FCS [Gibco], 1% Penicillin/Streptomycin [Gibco], and 1% GlutaMAX [Gibco]) was added dropwise to a total volume of 50 mL, with a 1-minute incubation and mixing step after each doubling of the volume. Cells were centrifuged at 400 g for 5 minutes at 4°C, and the supernatant was discarded. The cell pellets were resuspended in 1 mL of PBS and counted using a CellDrop (DeNovix).

### Cell Culture and Drug Screening

PBMCs were adjusted to a final concentration of 2 × 10^6^ cells/mL in cell culture medium, with or without the addition of 750 ng/mL PHA-L (Roche) for cellular activation. A 100 μL aliquot of the cell suspension was seeded into 96-well plates. Compounds or DMSO controls were diluted twofold in cell culture medium, and 100 μL of the prepared solutions was added to the wells. Compound concentrations were used according to table 1. DMSO concentration was adjusted to compound concentrations. The PBMCs were cultured at 37°C, 5% CO_2_, and 95% relative humidity for 22 hours.

**Table 1:**
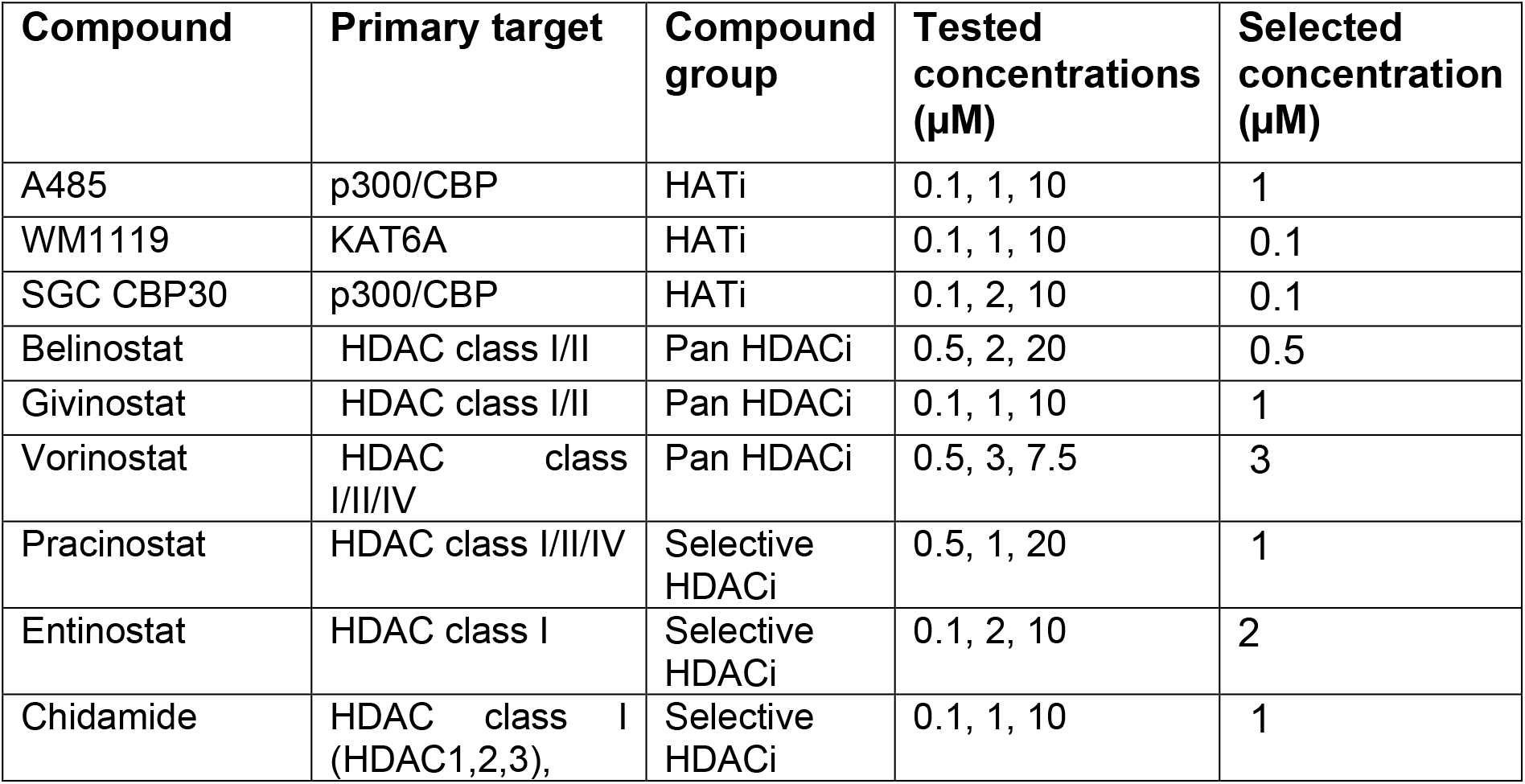
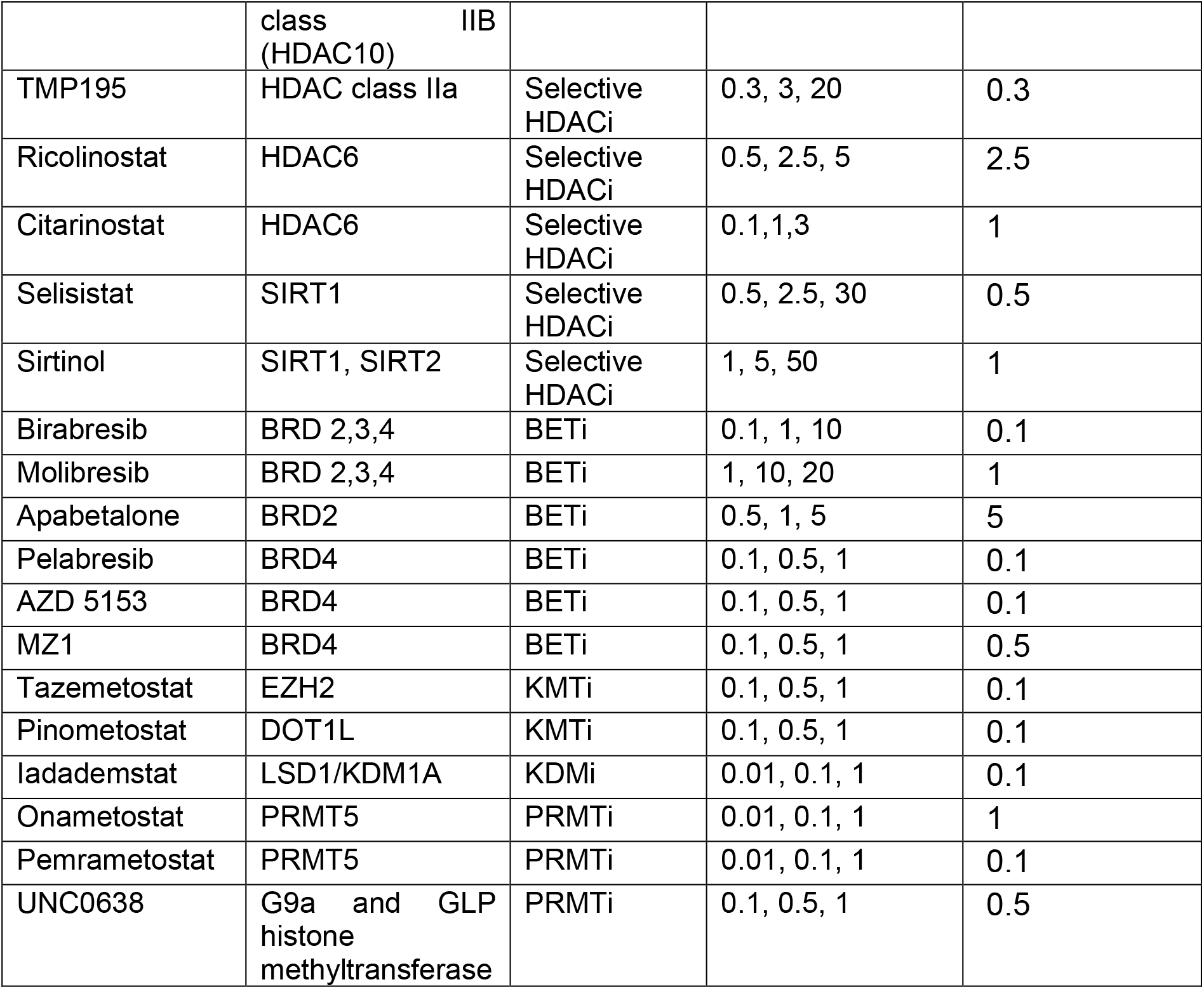
Epigenetic Inhibitor library, Compound names, primary targets, compound groups, tested concentrations (low, medium, high) and selected concentrations.

| Compound | Primary target | Compound group | Tested concentrations ($\mu\text{M}$) | Selected concentration ($\mu\text{M}$) |
| --- | --- | --- | --- | --- |
| A485 | p300/CBP | HATi | 0.1, 1, 10 | 1 |
| WM1119 | KAT6A | HATi | 0.1, 1, 10 | 0.1 |
| SGC CBP30 | p300/CBP | HATi | 0.1, 2, 10 | 0.1 |
| Belinostat | HDAC class I/II | Pan HDACi | 0.5, 2, 20 | 0.5 |
| Givinostat | HDAC class I/II | Pan HDACi | 0.1, 1, 10 | 1 |
| Vorinostat | HDAC class I/II/IV | Pan HDACi | 0.5, 3, 7.5 | 3 |
| Pracinostat | HDAC class I/II/IV | Selective HDACi | 0.5, 1, 20 | 1 |
| Entinostat | HDAC class I | Selective HDACi | 0.1, 2, 10 | 2 |
| Chidamide | HDAC class I (HDAC1,2,3), class IIB<br>(HDAC10) | Selective HDACi | 0.1, 1, 10 | 1 |
| TMP195 | HDAC class IIa | Selective HDACi | 0.3, 3, 20 | 0.3 |
| Ricolinostat | HDAC6 | Selective HDACi | 0.5, 2.5, 5 | 2.5 |
| Citarinostat | HDAC6 | Selective HDACi | 0.1, 1, 3 | 1 |
| Selisistat | SIRT1 | Selective HDACi | 0.5, 2.5, 30 | 0.5 |
| Sirtinol | SIRT1, SIRT2 | Selective HDACi | 1, 5, 50 | 1 |
| Birabresib | BRD 2,3,4 | BETi | 0.1, 1, 10 | 0.1 |
| Molibresib | BRD 2,3,4 | BETi | 1, 10, 20 | 1 |
| Apabetalone | BRD2 | BETi | 0.5, 1, 5 | 5 |
| Pelabresib | BRD4 | BETi | 0.1, 0.5, 1 | 0.1 |
| AZD 5153 | BRD4 | BETi | 0.1, 0.5, 1 | 0.1 |
| MZ1 | BRD4 | BETi | 0.1, 0.5, 1 | 0.5 |
| Tazemetostat | EZH2 | KMTi | 0.1, 0.5, 1 | 0.1 |
| Pinometostat | DOT1L | KMTi | 0.1, 0.5, 1 | 0.1 |
| Iadademstat | LSD1/KDM1A | KDMi | 0.01, 0.1, 1 | 0.1 |
| Onametostat | PRMT5 | PRMTi | 0.01, 0.1, 1 | 1 |
| Pemrametostat | PRMT5 | PRMTi | 0.01, 0.1, 1 | 0.1 |
| UNC0638 | G9a and GLP histone methyltransferase | PRMTi | 0.1, 0.5, 1 | 0.5 |

### Inhibitor Library and tested concentrations

### Flow Cytometry

*We used a* spectral flow cytometer (Cytek Aurora), equipped with a 3-laser configuration (405 nm, 488 nm, and 640 nm). Reagent concentrations and staining procedure were established as previously demonstrated by our group (21).

After incubation, the samples were placed on ice for 10 minutes, and cells were harvested directly into 1.4 mL FACS tubes. To prevent monocyte attachment, the wells were rinsed with PBS containing 10 mM EDTA (Gibco). Cells were centrifuged at 400g, 5min and resuspended in 25 μL PBS. Prior to antibody staining, 1 μL Human TruStain FcX™ (BioLegent) per sample was added, carefully mixed, and cells were incubated for 10 min on ice. 25 μL of Live-dead dye (Live Dead Aqua, ThermoFisher) was added in a 1:250 dilution and incubated with for 10 min on ice. Antibody cocktail was added according to table 2. Cells stained for 30 minutes at 4°C in the dark. Staining was stopped by adding 1 mL of PBS with 2% FCS. Samples were centrifuged at 400 g, 5 min. and resuspended in 75 μl PBS with 2% FCS.

**Table 2:** Antibodies used for flow cytometry (Fluorophore/marker combinations).

| **Panel 1** |  | **Panel 2** |  |
| --- | --- | --- | --- |
| Fluorophore | Surface marker | Fluorophore | Surface marker |
| BV421 | CCR7 | eFlour450 | HLA-DR |
| BV480 | TCR$\gamma\delta$ | BV570 | CD16 |
| BV570 | CD8 | BV650 | CD56 |
| BV650 | CD28 | BV711 | CD95 |
| FITC | CD69 | BV785 | PD-1 |
| PerCP | CD45RA | FITC | CD69 |
| PE-Cy7 | CD4 | PerCp-Cy5.5 | CD14 |
| APC | CD25 | PE-Cy7 | CD3 |
| APC-Cy7 | CD3 | APC-Cy7 | CD19 |
|  |  | APC-Fire810 | CD38 |

The samples were analyzed using the Cytek Aurora following a quality assurance step conducted with Cytek SpectroFlo® QC beads. To minimize interference from debris, the forward scatter threshold was set to a minimum of 50000 units. Data acquisition was performed at a rate of fewer than 6000 events per second, with a total of 250000 events collected per sample tube.

After the measurement, the experimental file was “live unmixed” in the SpectroFlo^®^ software. Compensation was applied with a minimum of 5%. FCS files were exported for further analysis.

### Data analysis of flow cytometry data

Data clean-up was performed using FlowAI software in R, to remove anomalies from the files concerning stability of the flow rate, the signal acquisition and the dynamic range (22). Files were imported into Flow Jo™ (version 10). Following data cleanup, all samples were manually pre-gated to remove remaining aggregates, debris, and doublets by evaluating the scatter profiles **(Supplementary Figure 1A)**. Further manual gating for individual PBMC subsets, respectively, was performed in accordance with the literature (23, 24) and is displayed in **Supplementary Figure 1A**. From each specified subset, the percentages of the respective population and the median fluorescence intensity (MFI) of the individual activation marker expression in the depicted subsets were extracted and exported in .csv file format.

### Down-stream analysis of Flow Cytometry Data

Percentage values of live cells for PBMC subsets and the MFI values for activation/exhaustion markers were imported into R. Changes in cell subset abundances were calculated as the log2 fold change (log2FC) of the fraction for each value, subtracted by the log2FC of the fraction for the DMSO control. The fractions for each cell type within a sample were calculated by dividing the count of that cell type by the total count of all cell types in the same sample. This method aligns with the approach described in the R package scProportionTest (25). The activation/exhaustion marker expression was determined by dividing the marker MFI of the compound treated sample by the respective marker MFI of the DMSO control (FC = MFI_compound_/MFI_DMSO_)).

### Concentration finding

Viability data (percentage of single cells), were imported into R. Viability values were normalized by dividing them by the viability of the corresponding DMSO control. A threshold of 70% viability relative to the DMSO control was established as the initial concentration screening point. To assess the overall effect of compound treatment on cell activation, a summed absolute median fold-change (SFC) was calculated. Activation marker expression values were normalized to their corresponding DMSO controls, and fold-changes were computed. The median fold-change across three replicates was calculated for each marker, and the SFC was defined as the total sum of these median fold-changes for all measured markers across all cell types.

### Statistical analysis

Statistical testing was performed in R. For evaluation of flow cytometry results, statistical comparisons were done by paired t-test, comparing the percentage of live-cell values of individual replicates of every condition to the corresponding DMSO control. P-values were adjusted for multiple testing using Benjamini Hochberg correction. Significance was defined as adjusted p-value (*P < 0.05, **P < 0.01, and ***P < 0.001).

### Bulk-RNA Sequencing

After drug treatment with selected concentrations (table 1), 400.000 PBMCs were harvested and placed into v-bottom plates (Thermo Scientific). Additional 200.000 cells were used for a control flow cytometry staining (see *Flow Cytometry*). Cells were centrifuged at 400 g for 8 min. and resuspended in 150μl RLT-Buffer(RNeasy Mini Kit (Qiagen)) containing 0,001% ß-Mercaptoethanol. RNA was isolated using the RNeasy Mini Kit (Qiagen) according to the manufacturer’s protocol. RNA-Seq was performed using QuantSeq 3’ mRNA-SeqLibrary Prep Kit FWD for Illumina (Lexogen) according to the manufacturer’s protocol. The libraries were sequenced at the Biomedical Sequencing Facility (BSF) at the Research Center for Molecular Medicine of the Austrian Academy of Sciences (CeMM) using the Illumina HiSeq 3000/4000 platform.

### Bulk-RNA Sequencing analysis

The raw reads were aligned to the human reference genome GRCh38 (December 2013) and raw gene counts were generated using STAR (26) after an initial quality check using FastQC. The raw counts were normalized using the median of ratios method implemented in DESeq2 (27) and differential expression analysis was performed using the Wald test in DESeq2, with donor ID and corresponding control sample as covariates in the model. Genes were considered differentially expressed if they had an absolute log2 fold chang ≥ 1 and a false discovery rate (FDR)-adjusted p-value < 0.05. Principal component analysis (PCA) was performed using using PCAtools (10.18129/B9.bioc.PCAtools).

### Enrichment analysis

Differentially expressed genes were then functionally enriched using the enrichR package in R (28). We utlizied the “DisGeNET” database and selected the top 5 terms for every comparison. Hits were filtered for inflammatory diseases.

To provide further insights, enrichment terms were clustered and summarized using our recently developed tool SummArIzeR (https://doi.org/10.5281/zenodo.14627791), Briefly, the databases employed for enrichment included “BioPlanet_2019,” “GO_Biological_Process_2023,” and “Reactome_2022” from enrichR. For each database and comparison, the top 10 enriched terms were selected, with terms associated with fewer than five genes excluded.

Similarities between terms were calculated using the Jaccard similarity of their associated genes. Clusters were derived from the resulting similarity matrix using the Walktrap community detection algorithm. Edges below the similarity threshold of 0.07 (Figure 2C), 0.09 (Figure 2D, Supplementary Figure 2B, Figure 3B) were excluded from further analysis.

**Figure 1.**
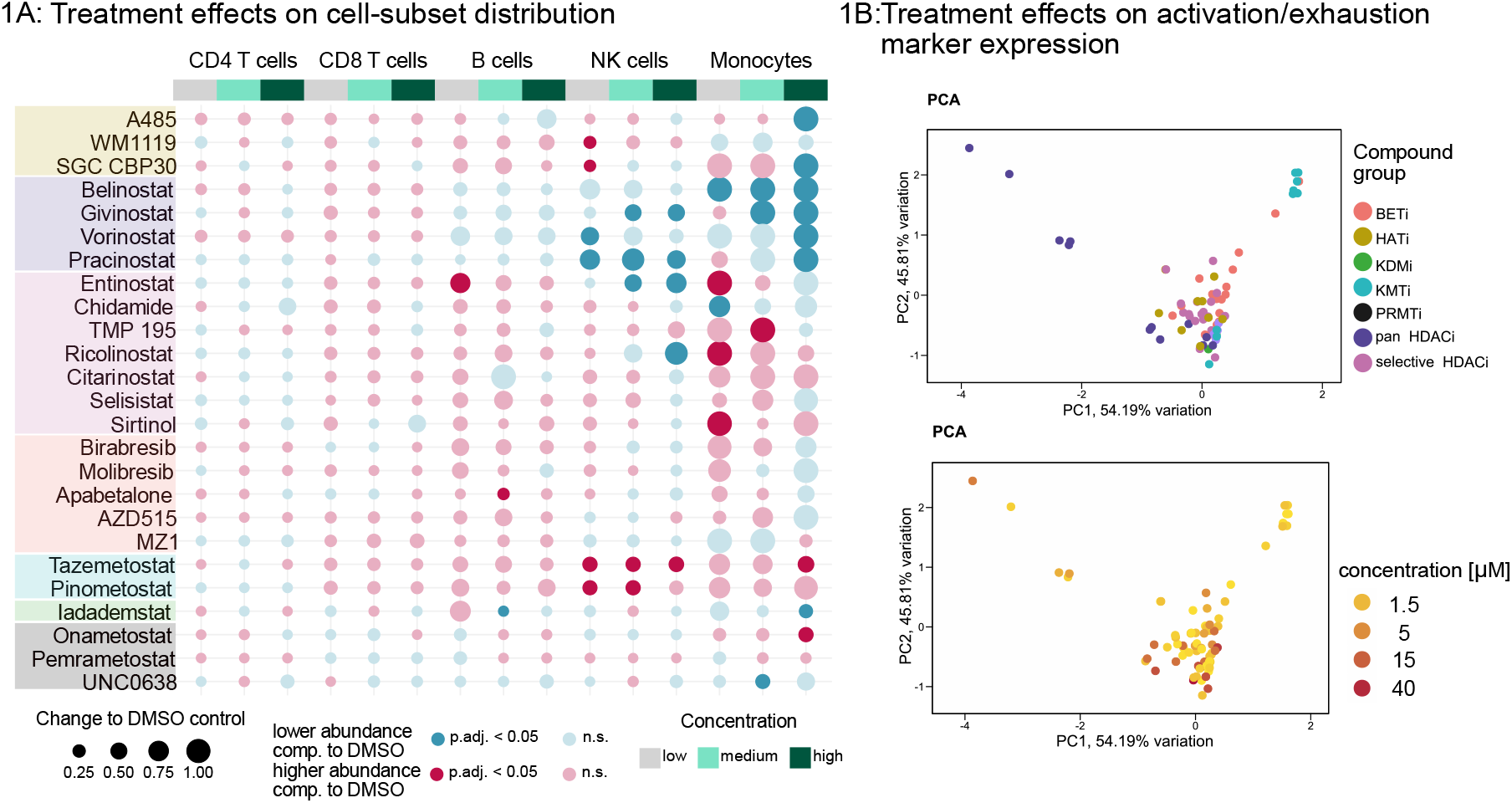
A. Bubble plot: Dot size depicting compound-induced perturbations in cell types, represented as changes relative to the DMSO control (Iog2(fraction of value)-log2(fraction of normvalue)), across 25 compounds and three concentrations (low, medium, high) (Table 1). Dot size reflects the magnitude of perturbations (capped at an absolute value of 1), while dot color indicates significance for up-or down-regulation (p-values calculated using paired t-test, comparing percentage of live cells for treated samples to DMSO control and adjusted with the Benjamini-Hochberg correction). C. Principal component analysis (PCA) of activation/exhaustion marker expression changes with color coding indicating compound groups. D. Principal component analysis (PCA) ofactivation/exhaustion marker expression changes with color coding indicating concentrations (in pM).

**Figure 2.**
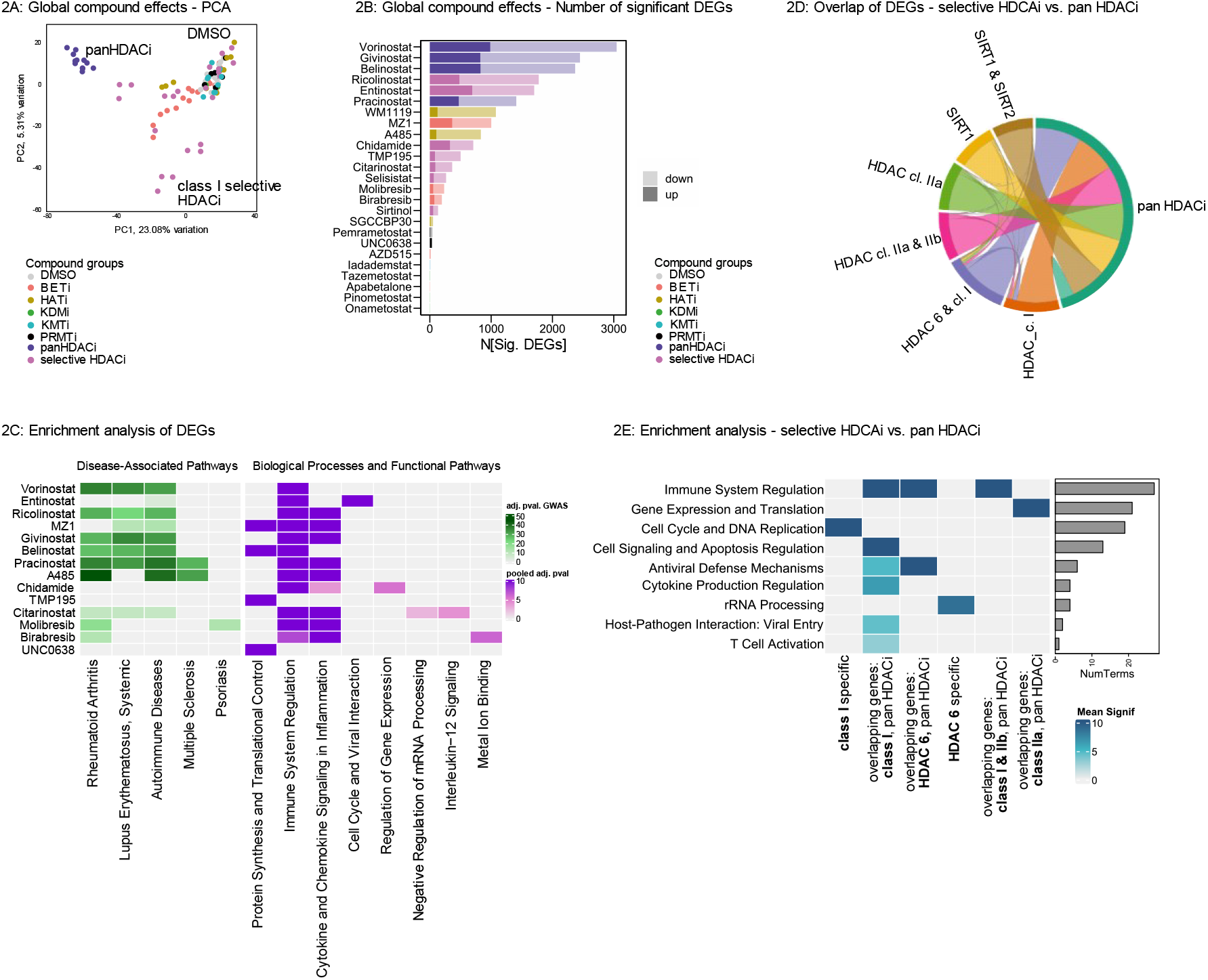
**A.**Principal component analysis (PCA) of bulk RNA sequencing data, displaying compound groups. Differential expression analysis: Differential expressed genes (DEGs) were calculated using DESeq2, (significant DEG (up-or downregulation)): baseMean > 5, |log2FoldChange| > 1, padj < 0.05)), overall 12732 genes with baseMean > 5 were detected. **B**. Stacked bar chart showing the number of significantly upregulated and downregulated genes compared to the DMSO control. **C**. Enrichment analysis of up-and downregulated DEGs between compound treatments and DMSO control. Left panel: Adjusted p-values (-log10(adj. pval.)) from the “DisGeNET” database filtered for inflammatory diseases. Right panel: Pooled adj. p-values (Fisher’s method (values capped at 10)) for clustered terms derived from “GO Biological Process 2023,” “Reactome 2022,” and “BioPlanet 2019” databases D. Chord plot depicting shared up– and downregulated DEGs (compound vs. DMSO) between selective and pan-HDAC inhibitors, stratification based on compound targets. F. Enrichment analysis of shared and unique up-and downregulated DEGs (compound vs. DMSO) for selective versus pan-HDAC inhibitors displaying adj. pooled p-values (Fisher’s method (values capped at 10)) for clustered terms derived from “GO Biological Process 2023,” “Reactome 2022,” and “BioPlanet 2019” databases.

Intersection of differentially expressed genes between compound targets was visualized using the chorddiag package in R (https://github.com/mattflor/chorddiag).

### Prediction of in-vivo drug effects

For comparison of drug target genes, we queried the Drug Gene Interaction Database (29) for 15 commonly used RA drugs and 15 non-inflammatory drugs.

RA drugs used: Methotrexate, Celecoxib, Rituximab, Leflunomide, Upadacitinib, Golimumab, Prednisone, Anakinra, Hydroxychloroquine, Abatacept, Baricitinib, Tocilizumab, Sulfasalazine, Tofacitinib, Certolizumab.

Non-inflammatory drugs used: Sildenafil, Taranabant, Vofopitant, Alprazolam, Raclopride, Tamsulosin, Ondansetron, Bupropion, Duloxetine, Lisinopril, Perhexiline Maleate, Tramadol, Lumateperone, Epinephrine Bitartrate, Temazepam, Pratosartan, Tadalafil, Idazoxan, Levocetirizine.

Associated pathways of target genes were analyzed doing an enrichment with EnrichR utilizing the TRRUST v2 database (30) and visualized in a network plot, Edges below the similarity threshold of 0.09 were excluded. Similarities between the associated pathways were calculated using the Jaccard similarity.

## Results

To explore the potential of epigenetic inhibitors for autoimmune disease treatment, we screened a library of 25 compounds targeting HATs, HDACs, BET proteins, PRMTs, KMTs, and KDMs, with varying degrees of target selectivity and different concentrations.

### Screening of Epigenetic Inhibitors Across Cell Subsets

HDACs play a pivotal role in various immune subsets. To screen the effects of epigenetic compounds on immune cell subsets we used spectral cytometry as an initial step. We could show pronounced effects of different compounds on cell-subset distributions, including NK cells, monocytes, and B cells, in a concentration-dependent manner **(Figure 1A, Supplementary Figure 1A)**.

The HAT Inhibitors A485 and WM1119 reduced monocyte levels at high concentrations, while WM1119 and SGC CBP30 upregulated NK cells at low concentrations. Pan-HDAC inhibitors had uniform effects across cell subsets, particularly reducing NK cells and monocytes. In contrast, selective HDAC inhibitors showed concentration-dependent effects, such as Entinostat upregulating monocytes at low concentrations but downregulating them at high concentrations. BET Inhibitors had minor effects on cell-subset distribution. The KMT Inhibitors Tazemetostat and Pinometostat increased NK cells and monocytes, while the demethylase inhibitor Iadademstat downregulated B cells and monocytes at distinct concentrations. PRMT Inhibitors showed fewer effects overall, with notable examples including Onametostat increasing monocytes and UNC0638 decreasing monocytes in a concentration-specific manner **(Figure 1A)**.

These findings revealed that compounds with similar targets share effects on the cell-subset distribution. Among the primary target cells of all tested compounds, we identified monocytes and NK cells.

### Activation and Exhaustion States of PBMC Subsets

To further assess the activation and exhaustion states of PBMCs, treated by epigenetic compounds, we analyzed the expression of surface activation and exhaustion markers such as CD69, CD25, CCR7, CD28 CD38, PD1, and CD95 **(Supplementary Figure 1B)**. Principal component analysis (PCA) of marker expression identified distinct clustering based on compound groups rather than concentrations, underscoring the role of target selectivity in shaping cellular responses **(Figures 1B, 1C)**. Pan-HDACi and HAT inhibitor treatments resulted in generally fewer activated cells, whereas selective HDAC inhibitors induced a more activated cellular phenotype **(Supplementary Figure 1B)**. For each compound, a single concentration for further analysis was selected by defining ≥70% cell viability compared to DMSO while maximizing changes in activation/exhaustion markers **(Supplementary Figure 1C)**.

### Transcriptomic Effects of Epigenetic Inhibitors

We performed bulk-RNA sequencing of PBMCs treated with the selected concentration for different epigenetic inhibitors (**graphical abstract**). The highest transcriptional changes were observed with pan-HDAC inhibitors followed by selective HDACi, HATi, and BETi **(Figure 2B)**. KMTi, KDMi, and PRMTi had minimal transcriptomic impacts, clustering closely with DMSO controls in the PCA **(Figure 2A, Figure 2B)**.

### Functional Enrichment of Differentially Expressed Genes

To investigate the biological function of differential expressed genes (DEGs), we performed an enrichment analysis. Interestingly, affected genes were associated with autoimmune diseases, in particular RA and systemic lupus erythematosus (SLE), for all pan-HDACi, the selective HDACi Ricolinostat and Citarinostat, the HATi A485, and the BETi Molibresib and Birabresib **(Figure 2C, left panel)**. Functional enrichment revealed immune system regulation as a dominant theme, alongside pathways such as translation, cytokine response, and viral interactions, in a compound-specific manner **(Figure 2C, right panel)**.

### Target-Specific and Selective Effects

To link the effects of the selective HDACi to their target, we performed a new stratification of the selective HDACi and calculated the intersection of differential expressed genes with the pan HDACi **(Figure 2D)**. We could indeed show that the inhibitors of the different targets are distinct from each other but overlap with the pan HDACi **(Figure 2D)** in their transcriptomic profiles. Enrichment analysis of overlapping genes highlighted that HDAC class I, HDAC6, and HDAC class I/IIb inhibitors were associated with “Immune System Regulation” pathways (Figure 2E). In contrast, overlapping genes between pan-HDACi and HDAC class IIa inhibitors were linked to terms associated with “Gene Expression and Translation.” Further analysis revealed that class I HDACi affected genes tied to “Cell Signaling and Apoptosis” as well as “Cell Cycle and DNA Replication.” Overlapping genes between pan– and HDAC6inhibitors were involved in “Antiviral Defense Mechanisms.” while HDAC6 inhibitor selective genes were enriched for pathways such as “rRNA Processing” (Figure 2E).

### RA-Specific Drug Effects

To validate findings in disease-relevant contexts, we compared *in-vitro* drug effects on PBMCs derived from HCs to RA patients. While overall responses were similar **(Figure 3A, Figure 3B)**, RA-specific differential expression emerged for genes associated with immune regulation, including Pracinostat-induced upregulation of cytokine signaling and Ricolinostat-induced downregulation of interferon signaling **(Figure 3C)**. We compared gene regulation patterns between HCs and DMSO-treated cells and those in RA patients and DMSO-treated cells. Ricolinostat and Vorinostat induced stronger treatment effects in RA samples, while Pracinostat showed enhanced regulation in HCs (Figure 3D). This trend was also evident in genes associated with “Immune System Regulation,” where Ricolinostat and Vorinostat demonstrated stronger regulation in RA, and Pracinostat exhibited more pronounced effects in HCs (Figure 3E). Overall, we could demonstrate similar drug effects for HCs and RA patients, with a RA selective downregulation of genes involved in “Immune System Regulation” for Ricolinostat.

**Figure 3.**
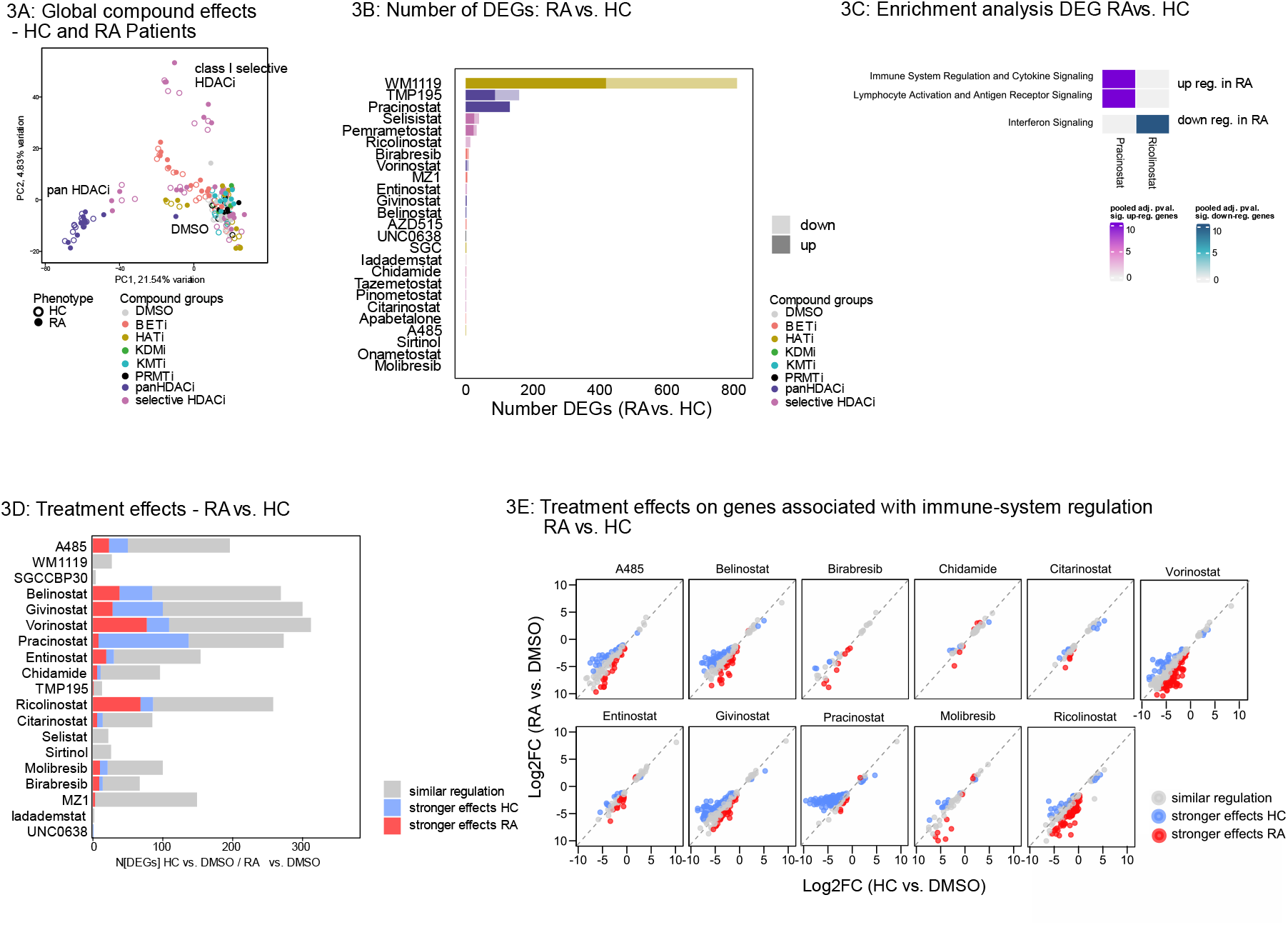
**A.**Principal component analysis (PCA) of bulk RNA sequencing data, displaying compound groups and patient characteristics (HC: healthy controls; RA: rheumatoid arthritis).Differential expression analysis: Differential expressed genes (DEGs) were calculated using DESeq2, (significant DEG (up-or downregulation)): baseMean > 5, |log2FoldChange| > 1, padj. < 0.05)), overall 12732 genes with baseMean > 5 were detected. **B**. Stacked bar chart showing the number of significantly upregulated and downregulated genes after compound treatment of samples from RA patients compared to healthy controls (HCs). **C**. Enrichment analysis for differentially expressed genes (RA vs. HC), up-and downregulation. The heatmap depicts pooled adj. p-values (calculated using Fisher’s method (values capped at 10)) for clustered terms derived from the “GO Biological Process 2023,” “Reactome 2022,” and “BioPlanet 2019” databases. **D**. Stacked bar chart showing effect ratios of DEGs (compound vs. DMSO). Effect ratios were calculated by comparing the absolute log2FoldChange ratios between HC and RA samples. Genes were classified as “stronger effects HC” if the ratio of | log2FoldChange _HC| to | log2FoldChange _RA| exceeded 1.25, indicating greater regulation in the HC condition and as “stronger effects RA” if the ratio of | log2FoldChange _RA| to | log2FoldChange _HC| exceeded 1.25, indicating greater regulation in the respective condition. Genes with ratios within this range were classified as “similar regulation.**E**. Scatterplot illustrating the l effect ratios of DEGs (compound vs. DMSO) of genes involved in “Immune-System Regulation” (from Figure 2C) in HCs and RA patients treated with different compounds.

### Prediction of in-vivo Effects

To model in vivo effects, we compared the gene targets of epigenetic inhibitors with those of established RA therapies and non-inflammatory drugs. Enrichment analysis revealed key TFs, including NFKB1, RELA, and STAT3, as shared regulatory nodes between conventional RA drugs and the epigenetic compounds **(Figure 4A)**. Jaccard similarity analysis of upstream TFs identified A485, Ricolinostat, and pan-HDAC inhibitors as having the strongest overlap with RA drugs, whereas Entinostat exhibited notably lower similarity **(Figure 4B)**.

**Figure 4.**
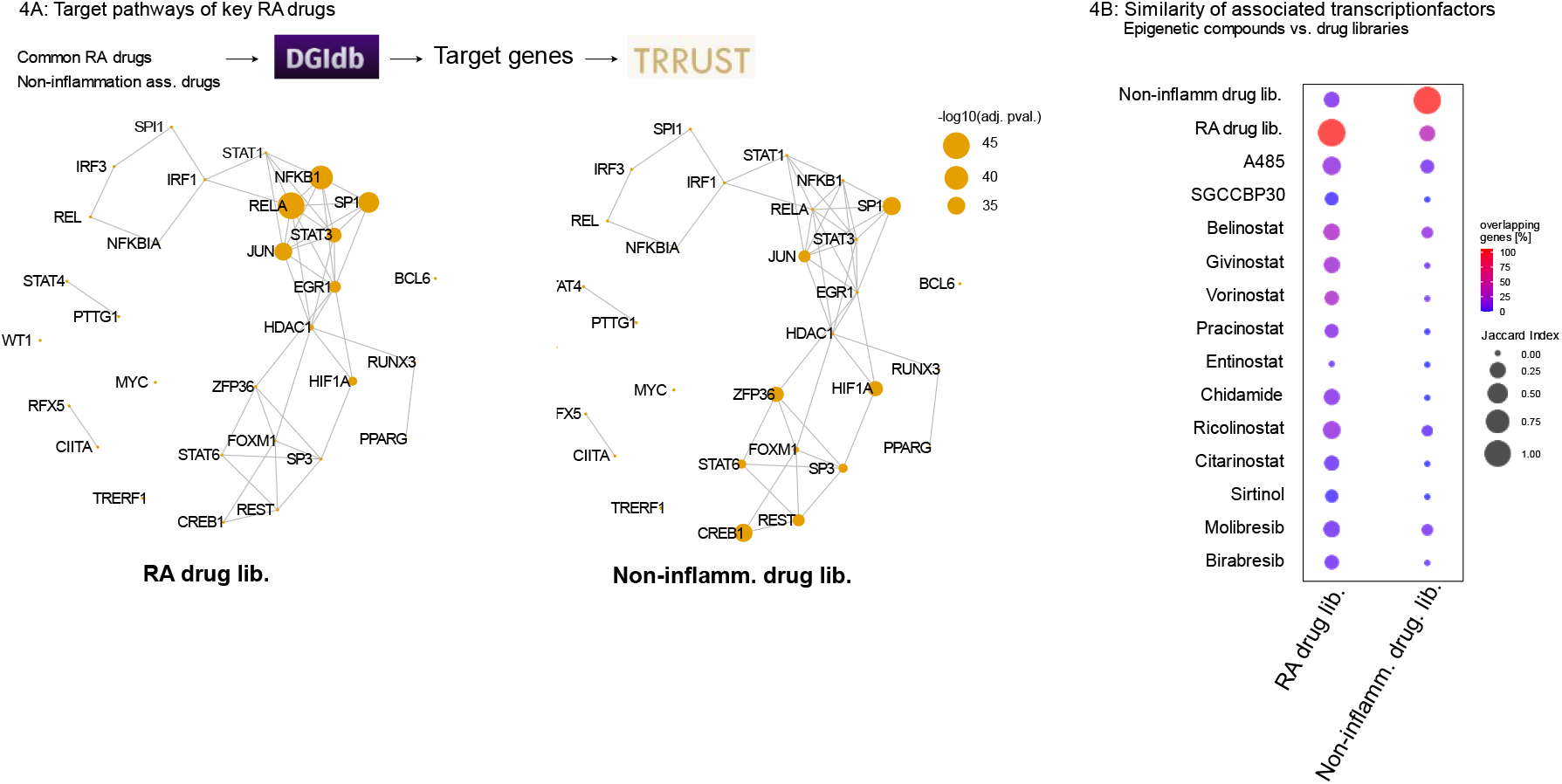
**A.**Associated transcription factors of 15 commonly used RA treatments and non-inflammatory drugs. Enrichment analysis of target genes was performed using the “TRRUST Transcription Factors 2019” database. The iGraph visualization shows associated transcription factors, with node size representing the adjusted p-value from enrichment analyses. **B**. Similarity of associated transcription factors between epigenetic compounds, 15 commonly used RA treatments and non-inflammatory drugs. Circle size depicting Jaccard Index, color depicting overlapping genes in percent. RA drugs used: Methotrexate, Celecoxib, Rituximab, Leflunomide, Upadacitinib, Golimumab, Prednisone, Anakinra, Hydroxychloroquine, Abatacept, Baricitinib, Tocilizumab, Sulfasalazine, Tofacitinib, Certolizumab.Non-inflammatory drugs used: Sildenafil, Taranabant, Vofopitant, Alprazolam, Raclopride, Tamsulosin, Ondansetron, Bupropion, Duloxetine, Lisinopril, Perhexiline Maleate, Tramadol, Lumateperone, Epinephrine Bitartrate, Temazepam, Pratosartan, Tadalafil, Idazoxan, Levocetirizine. Drug target genes were extracted from Drug Gene Interaction Database (29)

The presented results demonstrate that epigenetic inhibitors targeting histone acetylation modulate immune-related pathways in a target-specific manner, with RA-selective effects and overlaps with established drug pathways.

## Discussion

In this study, we demonstrated that epigenomic modulation of immune cells can reprogram pathogenic pathways in RA. Through a systematic screening of 25 inhibitors targeting histone acetylation and methylation, we identified distinct effects on immune cell subsets, activation states, and transcriptomic profiles. Key findings include concentration-dependent modulation of monocytes and NK cells, the regulation of inflammatory pathways through histone acetylation inhibitors, RA-specific gene regulation patterns, and overlap with transcriptional pathways targeted by established RA therapies.

Inhibitors of histone-modifying enzymes are successfully used in oncology but often cause adverse effects due to cytotoxic concentrations (31, 32). For autoimmune disease treatment, identifying more tolerable dosing strategies is crucial to minimize toxicity while maintaining therapeutic efficacy.

We screened three concentrations of a comprehensive epigenetic inhibitor library in PBMCs using flow cytometry, assessing cell subset distribution and activation as readouts. We observed concentration-dependent effects on cell-subset distribution and cell activation, with compound targets being the primary drivers of differential effects, while concentrations mainly influenced the effect size. Our findings suggests that beneficial compound effects can be preserved at lower concentrations which allowed us to select tolerable concentrations for the 25 screened compounds. In line with that, recent proteomic studies provided dose-response curves for multiple epigenetic inhibitors, showing that while acetylation levels stabilized beyond certain concentration thresholds, cell viability was reduced at higher concentrations (33).

Monocytes were the most altered PBMC type, with pan-HDACi and HATi reducing their abundance, and selective HDACi increasing their abundance. Monocytes play a critical role in inflammation by producing pro-inflammatory cytokines and presenting autoantigens (34, 35). In RA, monocytes infiltrate the synovium, where they can differentiate into pro-inflammatory macrophages or bone-resorbing osteoclasts (36, 37). Our findings identify monocytes as key targets of both HDACi and HATi. Consistently, HDAC1 and HDAC3 have been implicated in monocyte/macrophage differentiation (11). Moreover, both HDAC and BET inhibitors have been shown to reduce osteoclastogenesis *in-vitro*, further suggesting an impact on peripheral monocytes (17, 38, 39). However, whether monocytes become less abundant or rather downregulate characterizing surface marker, as described in studies with different *in-vitro* activation (40), needs to be further investigated.

Epigenetic inhibitors could already demonstrate anti-inflammatory effects *in-vivo* and *in-vitro*; however, uncovering global transcriptomic changes is essential to fully assess their potential beneficial outcomes and unintended adverse effects. To address this, we compared transcriptomic changes of multiple epigenetic inhibitors. We demonstrated transcriptomic changes primarily with acetylation-modifying enzyme inhibitors (HDACi, HATi, BETi), which were associated with autoimmune diseases such as RA and SLE. Most compounds in these groups targeted immune regulation pathways, consistent with previous studies (41, 42). Our data show target genes of compounds explaining known *in-vitro* effects. Other pathways included gene expression regulation and cell cycle control, which are known effects of epigenetic inhibitors from oncology research (43). Histone methylation-modifying enzymes exhibited minimal or no impact on the transcriptome. The limited transcriptional response to demethylase inhibition may reflect the relative stability of histone methylation marks compared to the more dynamic nature of acetylation (44).

The presented data revealed the potential of epigenetic drugs in regulating inflammation.

By screening a comprehensive inhibitor library with varying targets, we linked effects to specific epigenetic modifiers. Our data confirms HDAC class I as a key inflammatory regulator; however, targeting HDAC class I also affected apoptosis and cell cycle pathways, effects that were also previously described in oncology research (43). HDAC class I and IIb inhibition selectively enriched immune-related pathways to but not class IIa inhibition, which predominantly affected gene expression and translation. This discrepancy could be attributed to the fundamental differences in the roles and localization of class IIa and IIb HDACs (45).

Notably, HDAC6 emerged as a promising drug target, with selectively enriched immune-related pathways, consistent with recent *in-vitro* (46, 47) and *in-vivo* studies (48, 49). Unlike pan-HDAC inhibitors, HDAC6 inhibition appears to selectively impair rRNA processing without broadly affecting other transcriptional programs. The role of HDAC6 as a promising drug target was further reinforced by RA specific downregulation of Interferon signaling related genes following Ricolinostat treatment. By comparing upstream TFs with commonly used RA drugs, we demonstrated significant overlap of pan-HDACi, Ricolinostat. This suggests that *in-vitro* effects observed here could potentially translate into therapeutic success in RA.

Despite the promising findings, several limitations have to be considered. Currently available selective inhibitors still exhibit off-target effects by affecting multiple HDAC classes, which may obscure the precise contribution of individual targets. Further validation using a broader panel of highly selective inhibitors is necessary to confirm our observations and minimize potential off-target effects. Moreover, while our study identified immune regulatory effects and potential drug candidates, translation into clinical practice remains uncertain. Prospective validation through phase I/II clinical trials is required to assess the therapeutic efficacy and safety of these epigenetic modulators in patients with RA. Our data also indicate cell-type specific compound effects; however, as we only performed bulk sequencing, follow-up studies assessing transcriptomic changes at the cell-type level are essential.

Our data demonstrate that histone acetylation modifiers (HDACi, HATi, BETi) modulate immune cell types and immune regulatory pathways in a target-specific manner. Selective HDAC class IIb and HDAC6 inhibitors maintain immune regulatory features of pan-HDACi while reducing non-selective effects. HDAC6 inhibition also led to RA-specific downregulation of interferon signaling, highlighting its potential for RA treatment. The similarities of target pathways with successful RA treatment indicate the potential of epigenetic modulation as a novel treatment strategy.

## Data availability statement

Data will be made available on acceptance.

## Acknowledgment

We thank the “MedUni Wien Biobank” for their support in the storage and organization of patient samples. We also thank the Biomedical Sequencing Facility at CeMM for their assistance with next-generation sequencing and Thomas Krausgruber for guidance in the planning of sequencing experiments.

## References

1. Smolen JS, Aletaha D, Barton A, Burmester GR, Emery P, Firestein GS, et al. Rheumatoid arthritis. Nat Rev Dis Primers. 2018;4:18001.

2. Jang S, Kwon EJ, Lee JJ. Rheumatoid Arthritis: Pathogenic Roles of Diverse Immune Cells. Int J Mol Sci. 2022;23(2).

3. Di Matteo A, Bathon JM, Emery P. Rheumatoid arthritis. Lancet. 2023;402(10416):2019–33.

4. Mazzone R, Zwergel C, Artico M, Taurone S, Ralli M, Greco A, et al. The emerging role of epigenetics in human autoimmune disorders. Clin Epigenetics. 2019;11(1):34.

5. Nemtsova MV, Zaletaev DV, Bure IV, Mikhaylenko DS, Kuznetsova EB, Alekseeva EA, et al. Epigenetic Changes in the Pathogenesis of Rheumatoid Arthritis. Front Genet. 2019;10:570.

6. Zhang Y, Sun Z, Jia J, Du T, Zhang N, Tang Y, et al. Overview of Histone Modification. Adv Exp Med Biol. 2021;1283:1–16.

7. Ru J, Wang Y, Li Z, Wang J, Ren C, Zhang J. Technologies of targeting histone deacetylase in drug discovery: Current progress and emerging prospects. Eur J Med Chem. 2023;261:115800.

8. Barter MJ, Butcher A, Wang H, Tsompani D, Galler M, Rumsby EL, et al. HDAC6 regulates NF-kappaB signalling to control chondrocyte IL-1-induced MMP and inflammatory gene expression. Sci Rep. 2022;12(1):6640.

9. Hamminger P, Rica R, Ellmeier W. Histone deacetylases as targets in autoimmune and autoinflammatory diseases. Adv Immunol. 2020;147:1–59.

10. Ghizzoni M, Haisma HJ, Maarsingh H, Dekker FJ. Histone acetyltransferases are crucial regulators in NF-kappaB mediated inflammation. Drug Discov Today. 2011;16(11-12):504–11.

11. Tordera RM, Cortes-Erice M. Role of Histone Deacetylases in Monocyte Function in Health and Chronic Inflammatory Diseases. Rev Physiol Biochem Pharmacol. 2021;180:1–47.

12. Ellmeier W, Seiser C. Histone deacetylase function in CD4(+) T cells. Nat Rev Immunol. 2018;18(10):617–34.

13. Goschl L, Preglej T, Boucheron N, Saferding V, Muller L, Platzer A, et al. Histone deacetylase 1 (HDAC1): A key player of T cell-mediated arthritis. J Autoimmun. 2020;108:102379.

14. Zhang S, Meng Y, Zhou L, Qiu L, Wang H, Su D, et al. Targeting epigenetic regulators for inflammation: Mechanisms and intervention therapy. MedComm (2020). 2022;3(4):e173.

15. Xu WD, Huang Q, Huang AF. Emerging role of EZH2 in rheumatic diseases: A comprehensive review. Int J Rheum Dis. 2022;25(11):1230–8.

16. Subramanian S, Bates SE, Wright JJ, Espinoza-Delgado I, Piekarz RL. Clinical Toxicities of Histone Deacetylase Inhibitors. Pharmaceuticals (Basel). 2010;3(9):2751–67.

17. Friscic J, Reinwald C, Bottcher M, Houtman M, Euler M, Chen X, et al. Reset of Inflammatory Priming of Joint Tissue and Reduction of the Severity of Arthritis Flares by Bromodomain Inhibition. Arthritis Rheumatol. 2023;75(4):517–32.

18. Oh BR, Suh DH, Bae D, Ha N, Choi YI, Yoo HJ, et al. Therapeutic effect of a novel histone deacetylase 6 inhibitor, CKD-L, on collagen-induced arthritis in vivo and regulatory T cells in rheumatoid arthritis in vitro. Arthritis Res Ther. 2017;19(1):154.

19. Lin HS, Hu CY, Chan HY, Liew YY, Huang HP, Lepescheux L, et al. Anti-rheumatic activities of histone deacetylase (HDAC) inhibitors in vivo in collagen-induced arthritis in rodents. Br J Pharmacol. 2007;150(7):862–72.

20. Vojinovic J, Damjanov N. HDAC inhibition in rheumatoid arthritis and juvenile idiopathic arthritis. Mol Med. 2011;17(5-6):397–403.

21. Preglej T, Brinkmann M, Steiner G, Aletaha D, Goschl L, Bonelli M. Advanced immunophenotyping: A powerful tool for immune profiling, drug screening, and a personalized treatment approach. Front Immunol. 2023;14:1096096.

22. Monaco G, Chen H, Poidinger M, Chen J, de Magalhaes JP, Larbi A. flowAI: automatic and interactive anomaly discerning tools for flow cytometry data. Bioinformatics. 2016;32(16):2473–80.

23. Park LM, Lannigan J, Jaimes MC. OMIP-069: Forty-Color Full Spectrum Flow Cytometry Panel for Deep Immunophenotyping of Major Cell Subsets in Human Peripheral Blood. Cytometry A. 2020;97(10):1044–51.

24. Wang SR, Zhong N, Zhang XM, Zhao ZB, Balderas R, Li L, et al. OMIP 071: A 31-Parameter Flow Cytometry Panel for In-Depth Immunophenotyping of Human T-Cell Subsets Using Surface Markers. Cytometry A. 2021;99(3):273–7.

25. Miller SA, Policastro RA, Sriramkumar S, Lai T, Huntington TD, Ladaika CA, et al. LSD1 and Aberrant DNA Methylation Mediate Persistence of Enteroendocrine Progenitors That Support BRAF-Mutant Colorectal Cancer. Cancer Res. 2021;81(14):3791–805.

26. Dobin A, Davis CA, Schlesinger F, Drenkow J, Zaleski C, Jha S, et al. STAR: ultrafast universal RNA-seq aligner. Bioinformatics. 2013;29(1):15–21.

27. Love MI, Huber W, Anders S. Moderated estimation of fold change and dispersion for RNA-seq data with DESeq2. Genome Biol. 2014;15(12):550.

28. Kuleshov MV, Jones MR, Rouillard AD, Fernandez NF, Duan Q, Wang Z, et al. Enrichr: a comprehensive gene set enrichment analysis web server 2016 update. Nucleic Acids Res. 2016;44(W1):W90–7.

29. Cannon M, Stevenson J, Stahl K, Basu R, Coffman A, Kiwala S, et al. DGIdb 5.0: rebuilding the drug-gene interaction database for precision medicine and drug discovery platforms. Nucleic Acids Res. 2024;52(D1):D1227–D35.

30. Han H, Cho JW, Lee S, Yun A, Kim H, Bae D, et al. TRRUST v2: an expanded reference database of human and mouse transcriptional regulatory interactions. Nucleic Acids Res. 2018;46(D1):D380–D6.

31. Shi MQ, Xu Y, Fu X, Pan DS, Lu XP, Xiao Y, et al. Advances in targeting histone deacetylase for treatment of solid tumors. J Hematol Oncol. 2024;17(1):37.

32. Cheng B, Pan W, Xiao Y, Ding Z, Zhou Y, Fei X, et al. HDAC-targeting epigenetic modulators for cancer immunotherapy. Eur J Med Chem. 2024;265:116129.

33. Chang YC, Gnann C, Steimbach RR, Bayer FP, Lechner S, Sakhteman A, et al. Decrypting lysine deacetylase inhibitor action and protein modifications by dose-resolved proteomics. Cell Rep. 2024;43(6):114272.

34. Kawakami A, Iwamoto N, Fujio K. Editorial: The role of monocytes/macrophages in autoimmunity and autoinflammation. Front Immunol. 2022;13:1093430.

35. Yang S, Zhao M, Jia S. Macrophage: Key player in the pathogenesis of autoimmune diseases. Front Immunol. 2023;14:1080310.

36. Salnikova DI, Nikiforov NG, Postnov AY, Orekhov AN. Target Role of Monocytes as Key Cells of Innate Immunity in Rheumatoid Arthritis. Diseases. 2024;12(5).

37. McGarry T, Hanlon MM, Marzaioli V, Cunningham CC, Krishna V, Murray K, et al. Rheumatoid arthritis CD14(+) monocytes display metabolic and inflammatory dysfunction, a phenotype that precedes clinical manifestation of disease. Clin Transl Immunology. 2021;10(1):e1237.

38. Peng X, Wang T, Wang Q, Zhao Y, Xu H, Yang H, et al. Pan-histone deacetylase inhibitor vorinostat suppresses osteoclastic bone resorption through modulation of RANKL-evoked signaling and ameliorates ovariectomy-induced bone loss. Cell Commun Signal. 2024;22(1):160.

39. Park-Min KH, Lim E, Lee MJ, Park SH, Giannopoulou E, Yarilina A, et al. Inhibition of osteoclastogenesis and inflammatory bone resorption by targeting BET proteins and epigenetic regulation. Nat Commun. 2014;5:5418.

40. Bazil V, Strominger JL. Shedding as a mechanism of down-modulation of CD14 on stimulated human monocytes. J Immunol. 1991;147(5):1567–74.

41. Kim J, He Y, Tormen S, Kleindienst P, Ducoli L, Restivo G, et al. The p300/CBP Inhibitor A485 Normalizes Psoriatic Fibroblast Gene Expression In Vitro and Reduces Psoriasis-Like Skin Inflammation In Vivo. J Invest Dermatol. 2023;143(3):431–43 e19.

42. Bandukwala HS, Gagnon J, Togher S, Greenbaum JA, Lamperti ED, Parr NJ, et al. Selective inhibition of CD4+ T-cell cytokine production and autoimmunity by BET protein and c-Myc inhibitors. Proc Natl Acad Sci U S A. 2012;109(36):14532–7.

43. Ramaiah MJ, Tangutur AD, Manyam RR. Epigenetic modulation and understanding of HDAC inhibitors in cancer therapy. Life Sci. 2021;277:119504.

44. Chory EJ, Calarco JP, Hathaway NA, Bell O, Neel DS, Crabtree GR. Nucleosome Turnover Regulates Histone Methylation Patterns over the Genome. Mol Cell. 2019;73(1):61–72 e3.

45. Parra M. Class IIa HDACs – new insights into their functions in physiology and pathology. FEBS J. 2015;282(9):1736–44.

46. Park JK, Shon S, Yoo HJ, Suh DH, Bae D, Shin J, et al. Inhibition of histone deacetylase 6 suppresses inflammatory responses and invasiveness of fibroblast-like-synoviocytes in inflammatory arthritis. Arthritis Res Ther. 2021;23(1):177.

47. Lin D, Lai W, Zheng N, Luo H, Chen X, Que W, et al. Novel mechanistic study of HDAC6 regulation of rheumatoid arthritis via CMA: exploring potential therapeutic targets. Front Pharmacol. 2024;15:1383663.

48. Park JK, Jang YJ, Oh BR, Shin J, Bae D, Ha N, et al. Therapeutic potential of CKD-506, a novel selective histone deacetylase 6 inhibitor, in a murine model of rheumatoid arthritis. Arthritis Res Ther. 2020;22(1):176.

49. Yang HM, Lee C, Min J, Ha N, Bae D, Nam G, et al. Development of a tetrahydroindazolone-based HDAC6 inhibitor with in-vivo anti-arthritic activity. Bioorg Med Chem. 2024;99:117587.

