## Supplementary Figure 1 for "Targeting histone acetylation enables epigenetic modulation of inflammatory pathways – a novel therapeutic strategy for rheumatoid arthritis"

### Supplementary material

s1A: Gating Strategy

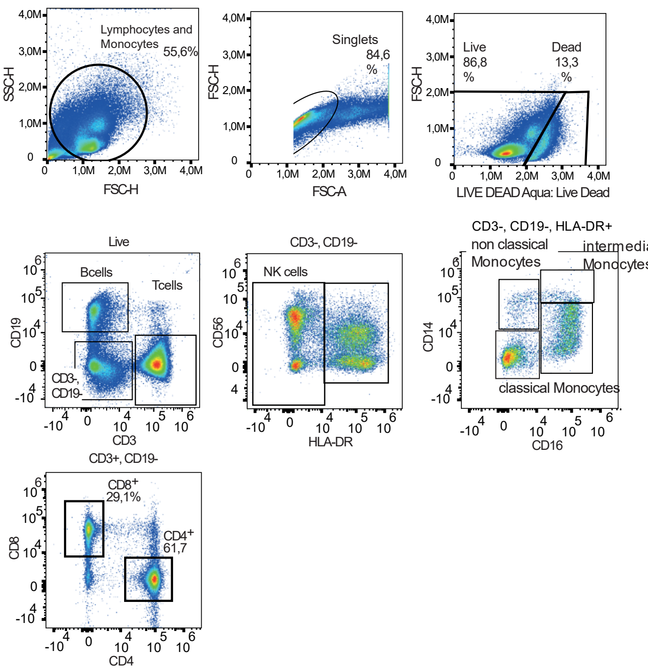

s1B: Activation marker expression

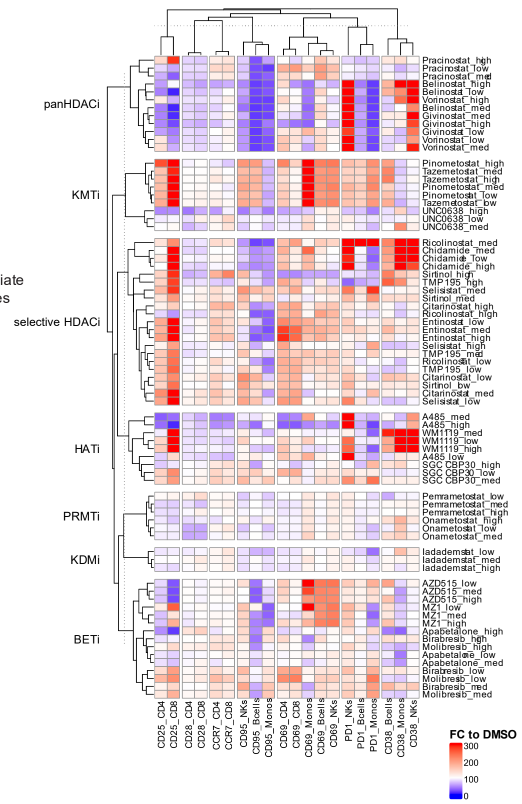

s1C: Concentration finding

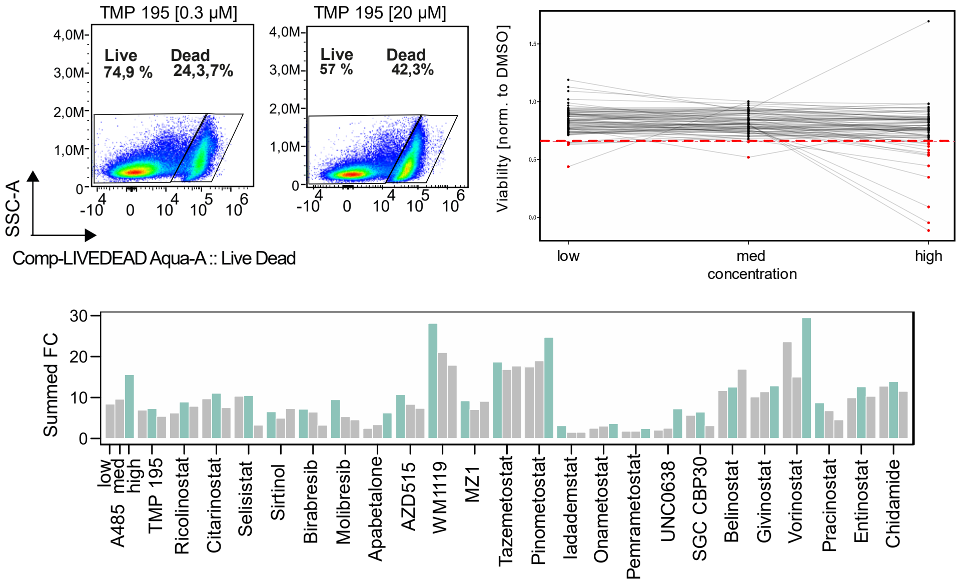

*Sup. Figure 1: A. Gating strategy employed for flow cytometry analysis. B. Heatmap displaying the fold change (FC) relative to DMSO for activation and exhaustion marker expression across the indicated cell types and compounds at three concentrations (FC = $MFI_{compound}/MFI_{DMSO}$). C. Example dot plot illustrating changes in cell viability for TMP*

*195 at the lowest and highest concentrations. The line plot depicts normalized viability (Viability norm. to DMSO = $Viability_{compound\text{-}treated}/Viability_{DMSO}$) with a red horizontal line indicating the 70% viability threshold used as the initial concentration screening point. Bar chart showing the summed absolute median fold-change (SFC). Activation marker expression values were normalized to their respective DMSO controls, and fold changes were calculated as in (B). The median fold change across three replicates was computed for each marker, and the SFC was defined as the total sum of these median fold changes ($SFC = \Sigma Median(FC_{marker})$). The maximum SFC for each compound is highlighted in green.*
